# cIAP1 inhibitor of apoptosis is a tumor suppressor in Ewing sarcoma

**DOI:** 10.1101/2025.06.27.661947

**Authors:** Florencia Cidre-Aranaz, Florian H. Geyer, Tilman L. B. Hölting, Tobias Faehling, Alina Ritter, Malenka Zimmermann, Lianghao Mao, Javier Alonso, Ana Sastre, Ashok Kumar Jayavelu, Maximilian M. L. Knott, Julian Musa, Thomas G. P. Grünewald

## Abstract

Ewing sarcoma (EwS) is a highly aggressive pediatric malignancy driven by EWSR1::ETS fusion oncoproteins –primarily EWSR1::FLI1– which deregulate genes essential for differentiation, proliferation, and cell survival. To uncover key downstream targets of this fusion involved in cell differentiation, we combined transcriptomic profiling of EwS cell lines following *EWSR1::FLI1* inhibition with gene ontology analysis, a clinically annotated gene expression dataset derived from EwS patient material and network analyses. This integrative approach identified inhibitor of apoptosis protein 1 (*cIAP1*, alias *BIRC2*) as an EWSR1::FLI1-suppresed gene. Despite its known oncogenic role in many cancers, *cIAP1* showed minimal expression in EwS. Using inducible *cIAP1* re-expression models in EwS cells, we demonstrated that *cIAP1* re-expression suppresses proliferation, clonogenic growth, and 3D spheroid formation in vitro. Transcriptomic and proteomic analyses revealed that low *cIAP1* expression enhances proliferation-related gene signatures, which are inhibited upon *cIAP1* re-expression. *In vivo* xenograft models revealed that *cIAP1* re-expression significantly reduces tumor growth, mitotic activity, and Ki-67 positivity, while increasing tumor necrosis and apoptosis. These findings highlight an unexpected tumor-suppressive role for *cIAP1* in fusion-driven sarcomas, contrasting with its pro-survival function in other cancers. Collectively, our results identify *cIAP1* as a prognostically relevant, EWSR1::FLI1-regulated hub whose re-expression disrupts tumor progression, offering a potential therapeutic strategy to restore tumor-suppressive pathways in EwS.

## MAIN TEXT

Ewing sarcoma (EwS) is an aggressive pediatric cancer with dismal overall survival rates for patients with relapsed or metastatic disease (<30% overall survival rates) (1). It is mainly characterized by a single genetic alteration: a chromosomal translocation fusing a member of the FET (*FUS, EWSR1, TAF15*) family of genes with an ETS transcription factor (mainly *FLI1* or *ERG*). The encoded FET:: ETS fusion proteins (in ∼85% of cases EWSR1::FLI1 and in ∼10% EWSR1::ERG) act as aberrant transcription factors that deregulate a plethora of genes involved in inhibition of cell differentiation, cell-cycle regulation, cell migration and proliferation among others (1).

To identify key *EWSR1::ETS* target genes we devised a systems biology approach where we first analyzed transcriptome profiles of eight EwS cell lines with or without shRNA-mediated knockdown (KD) of *EWSR1::FLI1* (KD <30% of baseline expression) for 96 h according to the workflow depicted in **Fig. 1a**. This analysis yielded a list of 1,318 differentially expressed genes (DEGs) being up- or downregulated (|log2 FC|≥0.75) after KD of *EWSR1::FLI1* across all cell lines. These DEGs were filtered for genes annotated with the gene ontology (GO) term ‘Regulation of Cell Differentiation’ using PantherDB, which was significantly enriched among the *EWSR1::ETS* regulated DEGs (*P*=8.77×10^−10^, false-discovery rate (FDR)=1.05×10^−7^). Using the obtained 167 DEGs (**Supplementary Table 1**), we carried out a network analysis using *Cytoscape* and the *GeneMania* plugin highlighting pathway, physical, and genetic interactions (2) (**Fig. 1b**). To identify key hubs within this network with potential clinical relevance, we further considered those genes whose expression significantly correlated with overall survival in a cohort of 196 EwS patients with matched gene expression and clinical data (*P*<0.01). This analysis resulted in 26 network hubs (**Supplementary Table 2**), out of which the single hub with the strongest correlation with patient overall survival was cellular inhibitor of apoptosis protein 1 (cIAP1; alias baculoviral IAP repeat containing 2, *BIRC2)* (**Fig. 1b,c**; *P*=0.0001).

**Fig. 1.**
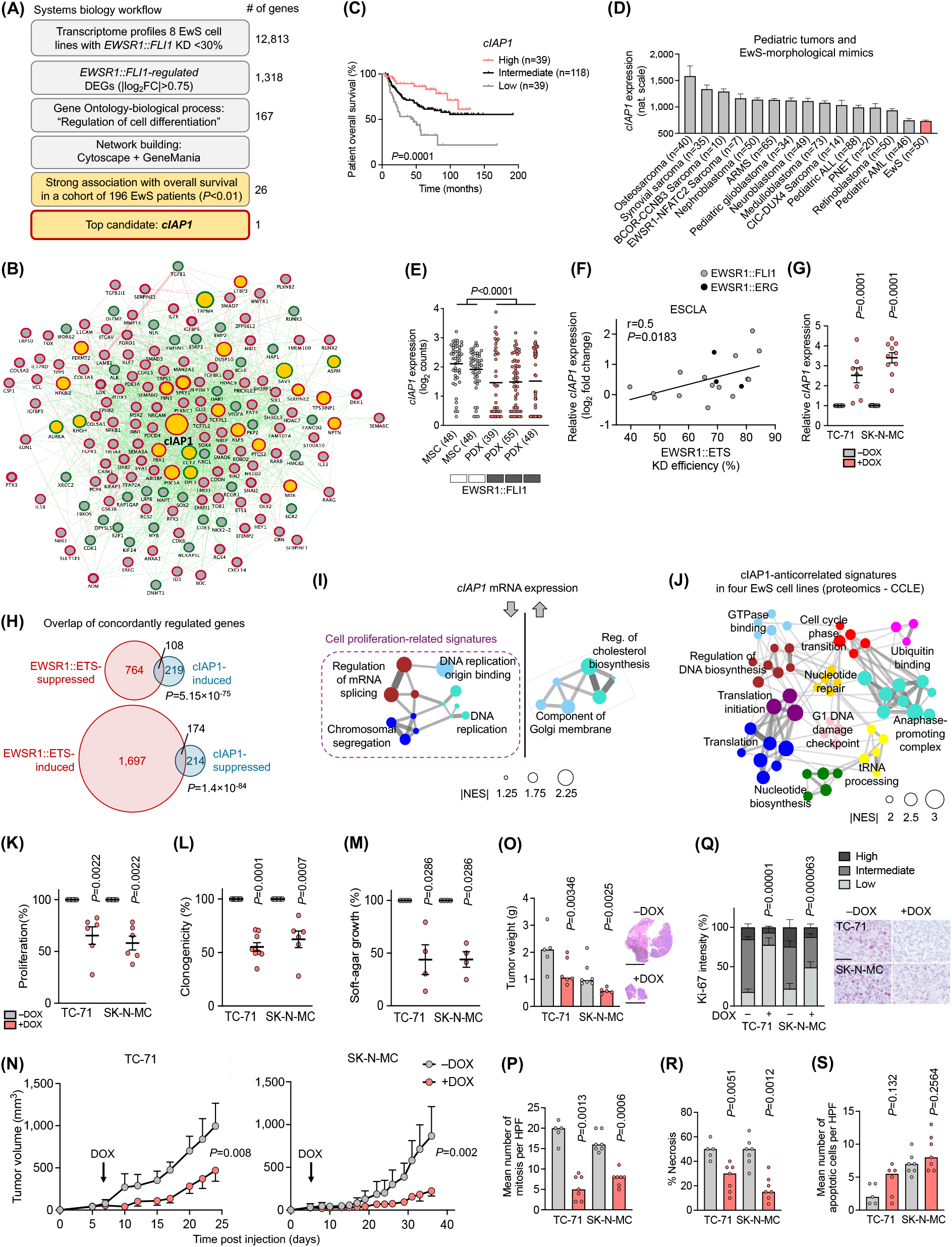
Systems biology approach identifies *cIAP1* as a prognostically relevant EWSR1::FLI1-regulated key hub displaying antitumorigenic functions in Ewing sarcoma. **a)** Workflow depicting a stepwise systems biology approach to identify DEGs regulated by EWSR1::FLI1, involved in regulation of cell differentiation, and associated with overall survival in a cohort of 196 EwS patients. Number of genes represent remaining candidates after each filtering step. **b)** Network of EWSR1::FLI1-regulated genes involved in regulation of cell differentiation. Genes are depicted as nodes (circles). Node outline color represent up-(green) or down-(red) regulation by EWSR1::FLI1. Node border width represent strength of regulation by EWSR1::FLI1 (thicker border represents higher the fold-change). Node size represents strength of association with overall survival in a 196 EwS patient cohort. Yellow nodes depict strong and significant association with overall survival (*P*<0.01). Connecting lines show three types of interconnections between nodes: physical (red), pathway (blue), genetic (green). **c)** Kaplan-Meier survival analysis of 196 primary EwS patients stratified by quintile *cIAP1* expression. Mantel-Haenszel test. DEG: differentially expressed genes; KD: knockdown. **d)** *cIAP1* expression levels of EwS and 14 additional primary pediatric tumors or EwS morphological mimics. Data are represented as bar plots where horizontal bars represent mean and SEM. The number of samples per group (*n*) is given in parentheses. ARMS: alveolar rhabdomyosarcoma; ALL: acute lymphocytic leukemia; PNET: primary neuroectodermal tumors; AML: acute myeloid leukemia. **e)** Analysis of single-cell RNA-Seq data (GSE XYZ) for *cIAP1* expression comparing EwS patient derived xenografts (PDX) to mesenchymal stem cells (MSC). Rectangles depict EWSR1::FLI1 expression level (white: low; black: high). Number of analyzed cells is given in parentheses. **f)** *cIAP1* and *EWSR1::ETS* expression in 18 EwS cell lines with conditional knockdown (KD) of *EWSR1::FLI1* (gray dots) or EWSR1::ERG (black dots) for 96 h measured by Affymetrix microarrays and qRT-PCR, respectively. *n*=*3* biologically independent experiments. **g)** Relative *cIAP1* expression as measured by qRT-PCR of TC-71 and SK-N-MC cells containing a DOX-inducible re-expression construct for *cIAP1. n*≥8 biologically independent experiments. Two-sided Mann-Whitney test. **h)** Size-proportional Venn diagrams of genes concordantly regulated 96 h after KD of *EWSR1::FLI1* or upregulation of *cIAP1* in TC-71 and SK-N-MC EwS cells. Minimum log_2_ fold-change ± 0.5. Fischer’s exact test. **i)** Weighted Gene Correlation Network Analysis (WGCNA) depicting functional gene enrichment of down- or up-regulated genes in *cIAP1* re-expressing EwS cells. Networks depict signatures presenting *P*<0.05, NES>1.25. NES, normalized enrichment score. Arrows depict direction of gene regulation. **j)** WGCNA of protein-sets obtained by Pearson correlation analysis of proteins whose expression correlate with cIAP1 protein expression in four EwS cell lines present in the proteomic dataset from the Cancer Cell Line Encyclopedia (CCLE). Networks depict signatures presenting *P*<0,05, NES>2. **k)** Viable cell count of TC-71 and SK-N-MC cells containing a DOX-inducible re-expression construct for *cIAP1* 72 h after treatment with or without DOX. Data are mean and SEM, *n*=6 biologically independent experiments. Two-sided Mann-Whitney test. **l)** Relative colony formation as measured in clonogenic index of TC-71 and SK-N-MC cells containing a DOX-inducible re-expression construct for *cIAP1*. Cells were grown either with or without DOX. *n*≥6 biologically independent experiments. Two-sided Mann-Whitney test. Data are mean and SEM. **m)** Relative percentage of area covered by colonies grown in soft-agar of TC-71 and SK-N-MC cells containing a DOX-inducible re-expression construct for *cIAP1*. Cells were grown either with or without DOX. *n*=4 biologically independent experiments. Two-sided Mann-Whitney test. **n)** Growth of EwS subcutaneous xenografts of TC-71 and SK-N-MC cells containing a DOX-inducible re-expression construct for *cIAP1* (arrow indicates start of DOX-treatment). Data are represented as means (*n*≥5 animals/group). Two-sided Mann-Whitney test. **o)** *Ex vivo* analysis of tumor weight (left) and representative H&E images of tumors from animals treated without or with DOX (right), scale bar = 5mm. **p)** *Ex vivo* analysis of mitotic index of xenografted TC-71 and SK-N-MC cell lines. Data are mean, *n*≥5 animals/group. **q)** *Ex vivo* analysis of Ki-67 positivity of xenografted TC-71 and SK-N-MC cell lines. Horizontal bars represent means and whiskers SEM, *n*≥5 animals/group. *P*-values were determined via χ^2^ test testing all positives (high and moderate immunoreactivity) versus negatives. Histological images depict representative Ki-67 micrographs. Scale bar=50μm. **r)** *Ex vivo* analysis of necrotic area of xenografted TC-71 and SK-N-MC cell lines upon *cIAP1* re-expression. Two-sided Mann-Whitney test. **s)** Graph depicts mean number of apoptotic cells per high power field (HPF) of *n*≥5 tumors per group analyzed.

Inhibitor of apoptosis proteins are frequently overexpressed in cancer and contribute to therapeutic resistance (3,4). To explore the specific expression pattern of cIAP1 in other cancers compared to EwS, we analyzed 15 cancer entities comprising pediatric tumors and EwS morphological mimics using well-curated microarray data from a prior study of our laboratory (5,6). Interestingly, EwS ranked as the tumor entity with the lowest *cIAP1* expression in primary tumors (**Fig. 1d**). To illuminate the underlying cause of this expression pattern, we re-analyzed scRNA-seq data from three EwS patient-derived xenografts (EWSR1::FLI1 expression) (7), and two primary cultures of mesenchymal stem cells –proposed cells of EwS origin (8)– negative for EWSR1::FLI1 (7). A comparative analysis showed a significantly lower *cIAP1* expression in the EwS-derived samples (**Fig. 1e**), suggesting a potential regulatory mechanism between *EWSR1::FLI1* and *cIAP1*. To explore this possibility, we reanalyzed the *cIAP1* locus in ChIP-seq data derived from the same eight EwS cell lines as in **Fig. 1a**. Inspection of the *cIAP1* promoter region revealed a specific EWSR1::FLI1 binding in this region in 7/8 cell lines analyzed (**Suppl. Fig. 1a**). This peak coincided with a H3K27ac mark that became specifically active upon inhibition of EWSR1::FLI1 in both A-673 and SK-N-MC cell lines, indicating a potential repressive activity of EWSR1::FLI1 on *cIAP1* (**Suppl. Fig. 1a**). Further correlation analyses on a published gene expression dataset comprising 18 EwS cell lines with shRNA-mediated KD of *EWSR1::FLI1/ERG* for 96 h (9) showed that stronger EWSR1::ETS KD significantly correlated with higher *cIAP1* expression (**Fig. 1f**, *P*=0.0183; *r*=0.5). Collectively, these data suggested a strong and likely direct repressive regulation of *cIAP1* by EWSR1::FLI1 and EWSR1::ERG.

To better understand the interplay between *cIAP1* and EWSR1::ETS, we generated cell line models containing a doxycycline (DOX)-inducible re-expression of *cIAP1* in two EwS cell lines (SK-N-MC and TC-71), which we found previously to display a high genomic and phenotypic stability contributing to reproducibility of experimental results (10). As shown in **Fig. 1g**, addition of DOX to the culture medium induced a median of 2.7–3.6-fold re-expression of *cIAP1* at the mRNA level. Consistent with the hypothesis that *cIAP1* may be a repressed EWSR1::ETS downstream effector, transcriptome profiling of two EwS cell lines after either KD of EWSR1::FLI1 or re-expression of *cIAP1* showed a highly significant (*P*=5.15×10^−75^ or *P*=1.4×10^−84^) overlap of concordant DEGs supporting the idea that cIAP1 acts as a regulatory downstream hub of the fusion (**Fig. 1h**).

To further investigate the consequences of *cIAP1* suppression in EwS, we performed transcriptome profiling coupled with gene-set enrichment and weighted gene correlation network analysis (WGCNA) of SK-N-MC and TC-71 EwS cells with/without conditional re-expression of *cIAP1*. These analyses demonstrated that low *cIAP1* levels promote the overrepresentation of gene-sets involved in cell proliferation-related signatures (**Fig. 1i**), which was completely abrogated by *cIAP1* re-expression (**Fig. 1i**). In agreement with these results, correlation analysis of publicly available proteomic data of four EwS cell lines (including TC-71 and SK-N-MC) showed a striking overrepresentation of proliferation-related signatures being anti-correlated with cIAP1 expression (**Fig. 1j**).

Although *cIAP1* overexpression is associated with promoting cancer cell survival and therapy resistance in carcinomas (4,11), its role in fusion-driven sarcomas is less clear. To explore the functional role(s) of *cIAP1* in EwS, we employed our DOX-inducible *cIAP1* re-expression models (SK-N-MC and TC-71). As shown in **Fig. 1k**, *cIAP1* re-expression significantly reduced (*P*=0.0022) proliferation in both cell lines (**Fig. 1k**), which was not observed in empty vector controls (12). Moreover, long-term re-expression of *cIAP1* significantly inhibited the capacity for clonogenic growth in two-dimensional (2D) cultures (**Fig. 1l**), and anchorage-independent spheroidal growth in 3D (**Fig. 1m**). Interestingly, when the same two EwS cell lines were engineered to conditionally re-express a *cIAP1* deletion mutant affecting their RING domain (ΔRING), which regulates its stability (13), the *cIAP1*-mediated inhibition on EwS clonogenic growth was reverted, suggesting that the RING domain is required to suppress the tumorigenic phenotype of EwS cells (**Suppl. Fig. 1b, c**).

To investigate whether the strong *in vitro* anti-tumorigenic role of cIAP1 could be recapitulated in an *in vivo* setting, TC-71 and SK-N-MC cells with conditional re-expression of *cIAP1* were tested in a pre-clinical xenotransplantation mouse model. For this, each cell line was subcutaneously injected in the right flanks of immunocompromised mice, and once tumors were palpable, DOX (treatment) or sucrose (control) were delivered via the drinking water. As depicted in **Fig. 1n**, cIAP1 re-expression significantly impaired local tumor growth in both EwS cell lines, which was mirrored by significant decrease in tumor weight (**Fig. 1o**). Ex vivo immunohistochemical (IHC) evaluation of the tumors showed that cIAP1 re-expression drastically decreased the mitotic index and Ki-67 immunopositivity (**Figs. 1p,q**). Strikingly, the xenografts showed a significant difference in tumor necrotic area (*P*=0.0051 or *P=*0.0012) and a tendency to presenting higher apoptotic indices in histological sections (**Fig. 1r,s**). Similar experiments performed with both cell lines transduced with empty control vectors exhibited no significant differences in these phenotypes upon DOX-treatment as demonstrated previously (12). These results indicated that EWSR1::ETS-meditated downregulation of *cIAP1* promotes clonogenic and anchorage-independent growth as well as tumorgenicity and suppression of cell death of EwS cells.

Collectively, our data show that *cIAP1* is a prognostically-relevant, EWSR1::ETS-regulated network hub in EwS, and that its re-expression severely impairs tumor fitness in vitro and in vivo.

## MATERIAL AND METHODS

### Provenience of cell lines and cell culture conditions

Human EwS SK-N-MC (CVCL_0530) and TC-71 (CVCL_2213) cell lines were provided by the German Collection of Microorganism and Cell Cultures (DSMZ). HEK293T (CVCL_0063) were purchased from American Type Culture Collection (ATCC). All cell lines were cultured in RPMI 1640 medium with stable glutamine (Biochrom, Germany) supplemented with 10% tetracycline-free fetal bovine serum (Sigma-Aldrich, Germany), 100 U/mL penicillin and 100 µg/mL streptomycin (Merck, Germany) at 37 °C with 5% CO_2_ in a humidified atmosphere. Cell lines were routinely tested for *Mycoplasma* contamination by nested PCR, and cell line identity was regularly verified by STR-profiling.

### RNA extraction, reverse transcription, and quantitative real-time polymerase chain reaction (qRT-PCR)

Total RNA was isolated using the NucleoSpin RNA kit (Macherey-Nagel, Germany). 1 µg of total RNA was reverse-transcribed using High-Capacity cDNA Reverse Transcription Kit (Applied Biosystems, USA). qRT-PCR reactions were performed using SYBR green Mastermix (Applied Biosystems) mixed with diluted cDNA (1:10) and 0.5 µM forward and reverse primer (total reaction volume 15 µl) on a BioRad CFX Connect instrument and analyzed using BioRad CFX Manager 3.1 software. Gene expression values were calculated using the 2^−(ΔΔCt)^ method(14) relative to the housekeeping gene *RPLP0* as internal control. The thermal conditions for qRT-PCR were as follows: heat activation at 95 °C for 2 min, DNA denaturation at 95 °C for 10 sec, and annealing and elongation at 60 °C for 20 sec (50 cycles), final denaturation at 95 °C for 30 sec. Oligonucleotides were purchased from MWG Eurofins Genomics (Germany) and are listed below:

*RPLP0* forward: 5’-GAAACTCTGCATTCTCGCTTC-3’

*RPLP0* reverse: 5’-GGTGTAATCCGTCTCCACAG-3’

*cIAP1* forward: 5’-TGTCAACTTCAGATACCACTGG-3’

*cIAP1* reverse: 5’-CACCAGGTCTCTATTAAAGCCC-3’

*EWSR1::FLI1* forward: 5’-GCCAAGCTCCAAGTCAATATAGC-3’

*EWSR1::FLI1* reverse: 5’-GAGGCCAGAATTCATGTTATTGC-3’

### *cIAP1* overexpression experiments

To re-express *cIAP1* at physiological levels, we assessed the baseline expression levels of *cIAP1* in 18 EwS cell lines for which whole-transcriptome data from human Affymetrix Clariom D arrays was available within a published Ewing Sarcoma Cell Line Atlas (ESCLA; triplicates per group per cell line **(9)**). As suitable models, we chose SK-N-MC and TC-71 cells as they exhibited the lowest baseline *cIAP1* expression among these cell lines and proceeded with cloning as described in (15). Briefly, *cIAP1* cDNA was PCR-amplified using a SKNMCshEWSR1::FLI1 sample treated with DOX as a template, and using AgeI- and NotI-restriction site containing primers (forward: 5’-ATTAACCGGTGCCACCATGCACAAAACTGCCTC-3’; reverse: 5’-TAATGCGGCCGCTTAAGAGAGAAATGTACGAAC-3’), before cloning it into the multiple cloning site of a modified pTP vector (16). *cIAP1* non-functional mutants were cloned from the same cDNA using touchdown-PCR. A RING-domain truncated mutant was generated by using a different reverse primer (5’-TAATGCGGCCGCTTAAGAGAGAAATGTACGAACAGTACCCTTGATTATACCAGTTCGTTCTTCTTGCAAC-3’) and a final Tm of 55 °C. The generated inserts were double restriction-digested with AgeI and NotI (NEB) and ligated into the pTP backbone (16) using T4 ligase (NEB). Positive clones were identified by colony PCR and cultured in 100 mL of LB Broth containing 100 µg/mL ampicillin. Plasmids were extracted and purified using a Midi-Prep Kit (Macherey-Nagel). The correct insertion of full-length *cIAP1* cDNA, or RING-domain deficient mutant was verified by Sanger sequencing (sequencing primers: forward 5’-ACGTATGTCGAGGTAGGCGT-3’; reverse 5’-TTCGTCTGACGTGGCAGC-3’). Lentiviral particles were generated in HEK293T cells and used for transduction of SK-N-MC and TC-71 EwS cells using polybrene (8 µg/mL). Transduced cells were selected with 0.5 µg/mL puromycin. Re-expression of *cIAP1* in SK-N-MC and TC-71 cells was achieved by addition of DOX (1 µg/mL) to the culture medium. Cells were single-cell cloned and specific clones were selected and tested for re-expression of *cIAP1*. Only the clones exhibiting re-expression levels comparable to the expression FCs after *EWSR1::FLI1* silencing (see above) were selected for functional assays. Cells transfected with the empty vector were used as additional controls as described previously (12).

### Transcriptome analyses

To assess the potential effect of *cIAP1* on gene expression in EwS cells, microarray analysis was performed. To this end, 1.2×10^4^ cells per well were seeded in 6-well plates and treated with 1μg/μl DOX for 72 h (DOX-refreshment after 48 h). Thereafter, total RNA was extracted with the ReliaPrep miRNA Cell and Tissue Miniprep System (Promega) and RNA quality was assessed with a Bioanalyzer. All samples had an RNA integrity number (RIN)>9 and were hybridized to Human Affymetrix Clariom D microarrays. Gene expression data were quantile normalized with Transcriptome Analysis Console (v4.0; Thermo Fisher Scientific) using the SST-RMA algorithm as previously described (17). Annotation of the data was performed using the Affymetrix library for Clariom D Array (version 2, *Homo sapiens*) on gene level. DEGs with consistent and significant FCs across cell lines were identified as follows: Normalized gene expression signals were log2 transformed. To avoid false discovery artifacts due to the detection of only minimally expressed genes, we excluded all genes with a lower expression value than that observed for *ERG* of the respective cell lines (log2 expression signal of 7.08 for SK-N-MC and 6.57 for TC-71), which is known to be virtually not expressed in EWSR1::FLI1 positive EwS cell lines (18). The FCs of both *cIAP1* re-expressing EwS cell lines were calculated for each cell line separately. Then the FCs in the *cIAP1* re-expressing samples were normalized to that of the empty control cells. Then both FCs were averaged to obtain the mean FC per gene across cell lines. DEGs were determined as having a log2 FC >0.5 or <–0.5, respectively.

### Gene-set enrichment analysis (GSEA)

To identify enriched gene-sets, genes were ranked by their expression FC between the groups DOX (−) and DOX (+). GSEA was performed using the FGSEA R package (v 3.6.3) based on Gene Ontology (GO) biological processes terms from MSigDB (c5.all.v7.0.symbols.gmt)(19). GO terms were filtered for statistical significance (adjusted *P*<0.05) and a normalized enrichment score |(NES)|>2 or |(NES)|>1.25 (10,000 permutations), depending on the dataset. In order to construct a network, the Weighted Gene Correlation Network Analysis R package (WGCNA R)(20) was used. Briefly, a binary matrix of GO-terms × genes (where 1 indicates the gene is present in the GO term and 0 indicates it is not) was created. Then, the Jaccard’s distance for all possible pairs was computed to create a symmetric GO adjacent matrix. Clusters of similar GO terms were identified using dynamicTreeCut algorithm, and the top 20 % highest edges were selected for visualization. The highest scoring node in each cluster was determined as the cluster label (rName). The obtained network and nodes files were fed into Cytoscape (v 3.8.0) for network design and visualization as previously described (21).

### Proliferation assays

For proliferation assays, 5–8×10^5^ EwS cells per well (depending on the cell line) were seeded in triplicates per group in 6-well plates and treated with 1 µg/mL DOX for 72 h. Thereafter, cells including their supernatant were harvested and counted using standardized hemocytometers (C-Chip, Biochrom) and the Trypan-Blue (Sigma-Aldrich) exclusion method as described in (22).

### Clonogenic growth assays

For clonogenic growth assays, *cIAP1* re-expressing EwS cells were seeded in triplicates at low density (2×10^3^ cells) per well in 12-well plates and grown for 9–11 d (depending on the cell line) with/without DOX-treatment (renewal of DOX or vehicle every 48 h). Thereafter, colonies were stained with crystal violet (Sigma-Aldrich) and colony numbers and areas were measured with the ImageJ Plugin *Colony area*. The clonogenicity index was calculated by multiplying the counted colonies with the corresponding colony area.

### Sphere formation assays in soft agar

For the analysis of anchorage-independent growth, *cIAP1* re-expressing EwS cells and respective controls were pre-treated with/without DOX for 48 h before seeding. A base of 2 mL of agar 1:1 with 2× DMEM medium was poured in wells of 6-well plates and left for 30–60 min to solidify. Then, 5×10^3^ cells per well were seeded in 500 µl in triplicates per condition (– /+DOX). Cells were kept in culture for 9–11 d and new medium (–/+DOX) was added on top of each well every 48 h. Spheres were stained with 50 µl of a 5 mg/mL MTT solution that was added dropwise to each well and incubated for 1h. Pictures of the stained spheres were taken, and their area was analyzed with ImageJ.

### Protein extraction and western blot

TC-71 and SK-NM-C EwS cell lines expressing DOX-inducible expression cassettes of *cIAP1* were treated with DOX (1 µg/mL) for 72 h. Whole-cell protein lysates were directly extracted using ice-cold RIPA buffer (SERVA Electrophoresis GmbH, Heidelberg, Germany) supplemented with 1X Halt™ Protease and Phosphatase Inhibitor Cocktail (Thermo Scientific) and further denatured and reduced by incubation in Laemmli SDS sample buffer (Thermo Scientific) at 95°C for 5 min. 20 µg of protein lysate per sample were separated on a 10% SDS-PAGE gel at 100 V and blotted on PVDF membranes using the Trans-Blot Turbo Transfer System (BioRad). Membranes were blocked for 1 h at RT in 5% skim milk (SERVA) in TBST (Carl Roth GmbH + Co. KG, Karlsruhe, Germany). Mouse monoclonal anti-cIAP1 antibody (1:500 sc-271419, Santa Cruz Biotechnology, Inc., Dallas, TX, USA) or rabbit monoclonal anti-GAPDH antibody (1:1,000, #2118, Cell Signaling Technology Europe B.V. Leiden, The Netherlands) were used as primary antibodies. Horseradish peroxidase (HRP) coupled anti-rabbit IgG (1:5,000, sc-2357, Santa Cruz) or anti-mouse IgG (1:5,000, A9044, Sigma-Aldrich) were used as secondary antibodies. Proteins bands were detected and visualized using chemiluminescence and Immobilon Western HRP Substrat (Sigma-Aldrich).

### *In vivo* experiments in mice

3×10^6^ SK-N-MC or TC-71 EwS cells harboring a re-expression construct for *cIAP1* were injected in a 1:1 mix of cells suspended in PBS with Geltrex Basement Membrane Mix (ThermoFisher) in the right flank of 10–12 weeks old NOD/scid/gamma (NSG) mice as described in(23). Tumor diameters were measured every second day with a caliper and tumor volume was calculated by the formula L×l^2^/2, where L is the length and l the width. When the tumors reached an average volume of 80 mm^3^, mice were randomized in two groups of which one was henceforth treated with 2 mg/mL DOX (Beladox, Bela-pharm, Germany) dissolved in drinking water containing 5% sucrose (Sigma-Aldrich) to induce an *in vivo* re-expression (DOX (+)), whereas the other group only received 5% sucrose (control, DOX (−)). Once tumors of control groups nearly reached an average volume of 1,500 mm^3^, all mice of the experiment were sacrificed by cervical dislocation. Other humane endpoints were determined as follows: Ulcerated tumors, loss of 20% body weight, constant curved or crouched body posture, bloody diarrhea or rectal prolapse, abnormal breathing, severe dehydration, visible abdominal distention, obese Body Condition Scores (BCS), apathy, and self-isolation. All tumor-bearing mice were sacrificed by cervical dislocation at the predefined experimental endpoint, when the mice reached a humane endpoint as listed above. After extraction of the tumors, they were weighted, and a small fraction of each tumor was snap frozen in liquid nitrogen to preserve the RNA isolation, while the remaining tumor tissue was fixed in 4 % formalin and embedded in paraffin for immunohistology. Animal experiments were approved by the governments of Upper Bavaria and conducted in accordance with ARRIVE guidelines, recommendations of the European Community (86/609/EEC), and United Kingdom Coordinating Committee on Cancer Research (UKCCCR) guidelines for the welfare and use of animals in cancer research.

### Survival analysis

Kaplan-Meier survival analyses were performed in 196 EwS patients (all samples derived from primary tumors that had been previously molecularly confirmed and retrospectively collected) that had been profiled at the mRNA level by gene expression microarrays in previous studies(24–27). Microarray data generated on Affymetrix HG-U133Plus2.0, Affymetrix HuEx-1.0-st or Amersham/GE Healthcare CodeLink microarrays of the EwS tumors (Gene Expression Omnibus (GEO) accession codes: <u>GSE63157</u> (24), <u>GSE12102</u> (25), <u>GSE17618</u> (26), <u>GSE34620</u> (27) provided with clinical annotations were normalized separately as previously described (5). Genes represented on all microarray platforms were kept for further analysis. Batch effects were removed using the ComBat algorithm (28). Data processing was done in R.

### Statistical analysis and software

Statistical data analysis was performed using PRISM 9 (GraphPad Software Inc., Ca, USA) on the raw data. If not specified otherwise in the figure legends, comparison of two groups in functional *in vitro* experiments was carried out using a two-sided Mann-Whitney test. If not specified otherwise in the figure legends, data are presented as dot plots with horizontal bars representing means and whiskers representing the standard error of the mean (SEM). Sample size for all *in vitro* experiments were chosen empirically. For *in vivo* experiments, the sample size was predetermined using power calculations with *β*=0.8 and *α*<0.05 based on preliminary data and in compliance with the 3R principles (replacement, reduction, refinement). Statistical differences between the groups were assessed by a Mantel-Haenszel test.

## Data availability

Original microarray data that support the findings of this study will be deposited at the National Center for Biotechnology Information (NCBI) GEO.

## ACKNOWLEDGEMENTS

We would like to thank Nadine Gmelin, Stefanie Kutschmann, Felina Zahnow, and Sabrina Knoth for their expert technical assistance, and Claudia Schmidt from the Light Microscopy Facility (German Cancer Research Center (DKFZ), Heidelberg, Germany) for her meticulous work in conducting immunohistochemical stainings.

## AUTHOR CONTRIBUTIONS

F.C.A and T.G.P.G. conceived the study. F.C.A. and T.G.P.G wrote the paper and drafted the figures and tables. F.C.A. carried out *in vitro* experiments and performed bioinformatic and statistical analyses. F.C.A., T.L.B.H. and J.M., performed *in vivo* experiments. F.H.G., A.R., M.M.L.K., T.F., M.Z., L.M. and A.K.J contributed to experimental procedures. A.S. and J.A. contributed patient data. All authors read and approved the final manuscript.

