## Supplementary figures and images for "cIAP1 inhibitor of apoptosis is a tumor suppressor in Ewing sarcoma"

### Supplementary Figure 1

#### Supplementary Figure S1

### (A)

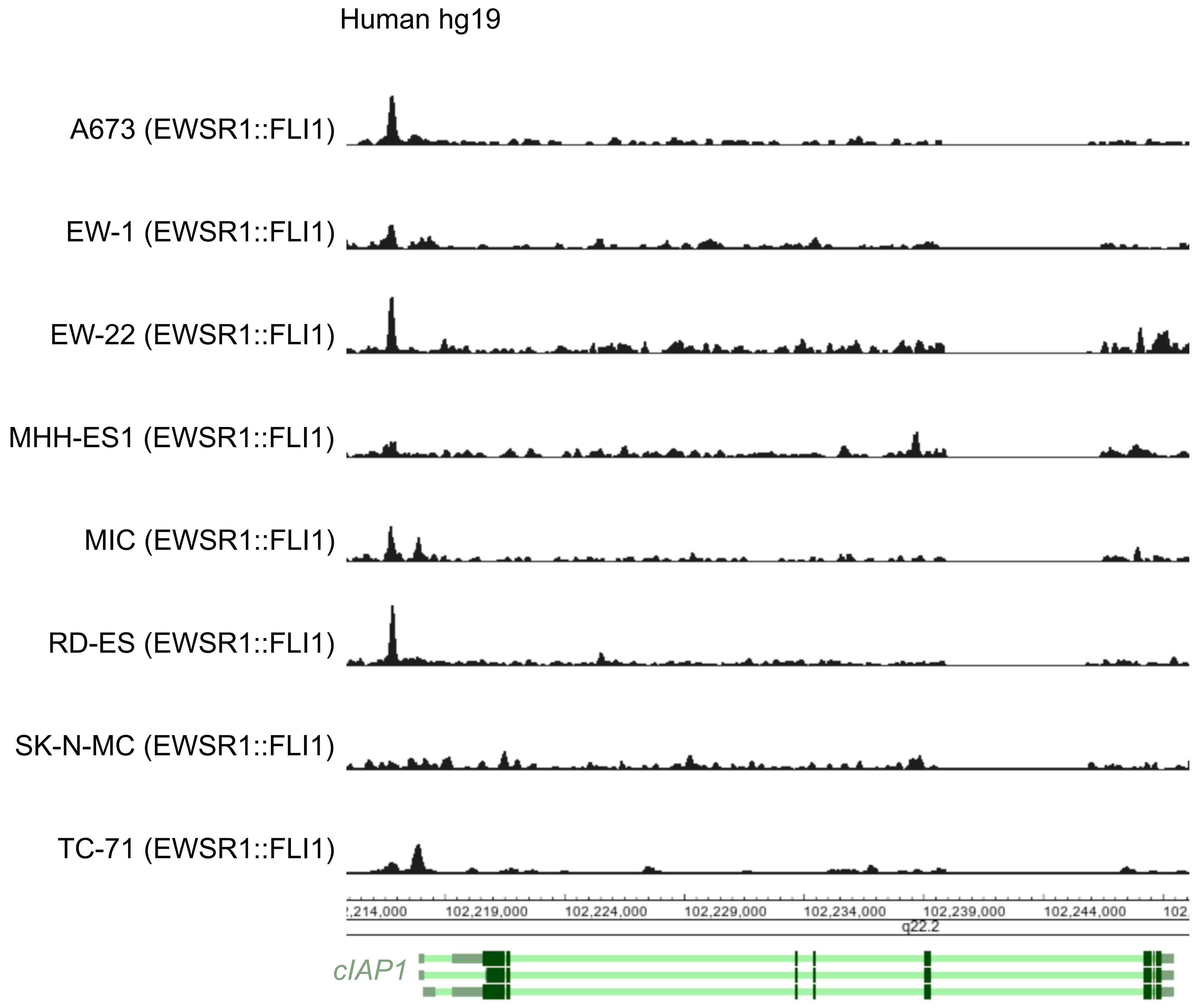


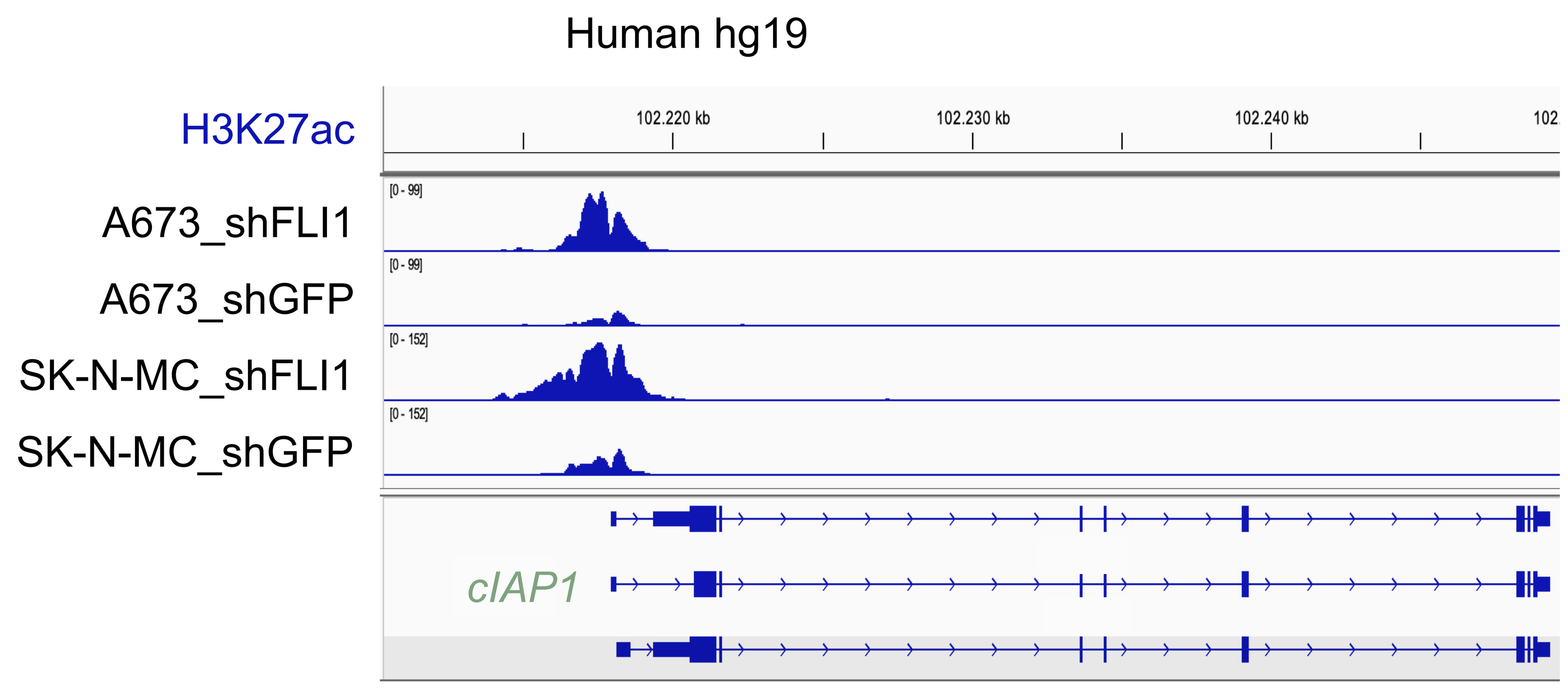


### (B)

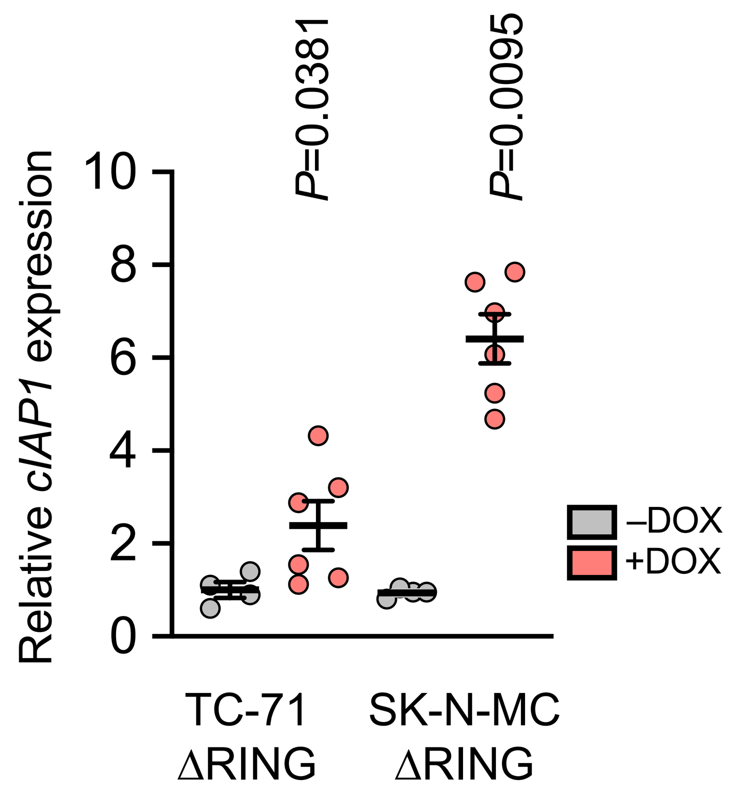


### (C)

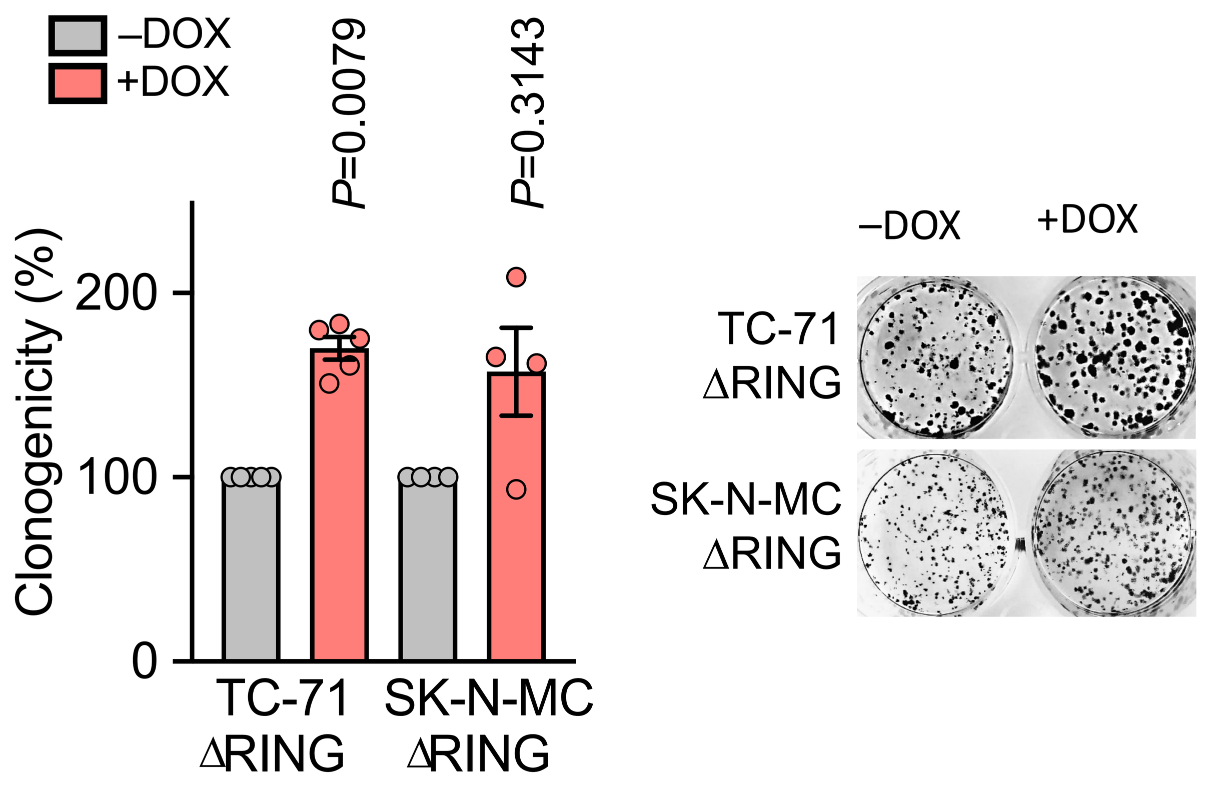
